# Anti-dengue viral activity of *Glycyrrhiza glabra* roots in Vero cells

**DOI:** 10.1101/2023.12.29.573598

**Authors:** Kalani Gayathri Jayasekera, Sugandhika Suresh, Charitha Goonasekera, Preethi Soyza, Namal Perera, Amitha Jayawardene, Kamani Gunasekera

## Abstract

Dengue is a global public health problem but no antiviral agent exists for dengue. Plants have been the source of many pharmaceutical agents. Therefore, we explored the antiviral potential of the aqueous extracts of *Glycyrrhiza glabra* (GG) roots on dengue viruses. Cell cytotoxicity assay with 3-(4, 5-dimethyl-2-thiazolyl)-2, 5-diphenyl-2H-tetrazolium bromide solution was used for estimating the maximum nontoxic dose (MNTD) and 50% cytotoxic concentration (CC_50_). Plaque reduction antiviral assay on Vero cells infected with dengue virus serotypes 1,2,3 and 4 were treated with dilutions of the MNTD of GG extract to calculate the half maximal inhibitory concentration (IC_50_). Selectivity index (SI) was calculated as CC_50_/IC_50_ ratio. Following bioactivity guided fractionation, MNTD, CC_50_, IC_50_ and SI values for fractions E and F were similarly obtained. Inhibition of NS1 antigen levels produced by dengue serotype-1 infected monocytes was studied in the presence and absence of GG extract after 48 hours. GG MNTD and CC_50_ were 166.7 µg/mL and 651.9 µg/mL. GG extract showed 98-100% inhibition of all four dengue serotypes at the MNTD and the IC_50_ values of the four serotypes ranged from 10-50 µg/mL indicating moderate pan-serotype inhibitory activity. Cytotoxicity of GG against Vero cells was low (CC_50_ >500 µg/mL). SI of GG extract for all four dengue serotypes were >10, indicating it had good potential as an antiviral agent. Fraction E and F had low to moderate inhibitory activity for all four serotypes and showed good potential as an antiviral agent only for dengue 4. This is the first report of significant anti-dengue viral activity of GG extract on all four dengue serotypes.

**Author Summary:** Dengue is a global public health problem but no antiviral agent exists for dengue. This study explores the antiviral potential of the aqueous extract of *Glycyrrhiza glabra* (GG) roots on dengue viruses. Virus inhibition in Vero cell lines infected with dengue virus serotypes 1-4 and treated with GG extract was demonstrated. GG extract was 98-100% inhibitory to all four serotypes and had moderate pan-serotype inhibitory activity. It had good potential as an antiviral agent with low cytotoxicity. This is the first report of significant anti-dengue viral activity of GG extract on all four dengue serotypes.

## Introduction

Dengue is a global public health problem. A recent estimate states that 390 million dengue infections occur per year, of which 96 million manifest clinically [1]. There are four serotypes of dengue virus in circulation. The enveloped single stranded positive sense RNA virus produces a single polyprotein, which is cleaved into three structural proteins (capsid, membrane and envelope) and seven non-structural proteins (NS1, NS2a, NS2B, NS3, NS4a, NS4B and NS5) that are associated with pathogenicity [2].

There are no specific antiviral agent for dengue at present. Mosquito control methods have so far been inadequate for containing spread, although *Wolbachia* infected mosquitoes seem to show promise for reducing the mosquito population [3]. The recently approved live-attenuated vaccine (Dengvaxia) is not recommended for seronegative populations because of the increased risk of severe dengue upon subsequent infection [4].

Clinical trials with repurposed pharmaceuticals have so far been disappointing [5]. A large reservoir of lead compounds potentially exists in nature, which could be used either directly or serve as lead structures on which new anti-dengue viral agents could be modelled [6]. Plants are the direct or indirect sources of approximately 60% of approved drugs and seven out of ten synthetic drugs are modelled on a natural product [7]. An anti-viral agent for dengue, is the need of the hour. Most studies have shown that severe dengue is more likely in those with high viral loads [8]. This has led to the belief that a drug that lowers viraemia by one to two orders of magnitude, may be associated with a favourable prognosis [9]. Antiviral drugs when given early would reduce the viral load and the duration of viraemia in dengue patients. This would not only prevent progression to severe disease but would also reduce the transmission from infected humans to mosquitoes.

Medicinal plants and their products have been used to cure fevers and other ailments in ancient traditional medicine. In Sri Lankan traditional medicine extracts from the whole plant of *Munronia pinnata* (MP), leaves of *Psidium guajava* (PG), dried flowers of *Aegle marmelos* (AM) and roots of *Glycyrrhiza glabra* are used for treating fever patients. We studied these plants but as reported previously MP, PG and AM did not adequately inhibit all four dengue serotypes and therefore, were not studied further [10].

The present study reports the pan-serotype inhibitory activity of aqueous extract of GG roots in Vero cells. GG is commonly known as liquorice and locally as Val mi [11]. It is a flowering perennial herb with a 0.5–1.5 m high stem including an elliptic leaflet. Bright yellow colour roots have a distinct sweet taste and it has a broad range of medicinal properties; both the roots and rhizomes are major medicinal parts of GG [11].

Bioassay guided fractionation of the aqueous extract of GG resulted in the isolation of fractions E and F, but only dengue serotypes 1 and 4 were significantly inhibited by these fractions. NS1 antigen has been found to be significantly associated with severe dengue infections [12]. Since aqueous extract of GG roots was found to be a better inhibitor of all four serotypes, further evaluation was carried out with dengue virus infected monocytes to determine its effects on NS1 antigen production.

## Methods

### Selection of plants

Out of 52 medicinal plants used traditionally to treat fever, four were shortlisted for bioassay-guided screening as, i) previously reported anti-dengue viral phytochemicals were present; ii) anti-viral effect against other viruses had been reported; and iii) they had been shown to be hepatoprotective. Based on these criteria, MP whole plant, GG roots, PG leaves and AM dried flowers were selected. The following description is limited to findings with aqueous extract of GG.

### Preparation of aqueous extract of GG roots

Healthy fresh GG roots were bought from Manning Market (an open market), Pettah in Sri Lanka and authenticated by the Bandaranayke Memorial Ayurveda Research Institute, Navinna, Sri Lanka (Registration no. 1603MS2017001). The GG roots were washed well in running tap water until the specimens were visibly clean. The washed roots were drained, air dried and cut into small pieces and washed in distilled water, followed by double distilled water. Sixty grams of GG was heated in 1920 mL of water in a beaker at 70°-80°C until reduced to 240 mL according to the traditional method of preparation. This procedure was repeated to get a total volume of 2 L of GG extract. The filtered extract was lyophilized and aliquots were stored at -20°C until used.

### Cells and viruses

African green monkey kidney (Vero) cells and *Aedes albopictus* larval cells (C6/36), obtained from the Dengue Research Centre, University of Sri Jayewardenepura, Sri Lanka, were grown in complete growth medium (CGM) that consisted of 12g/L of Dulbecco’s minimum essential medium (DMEM)/F12 (Sigma, USA D0547) supplemented with 5% fetal bovine serum (FBS) and 1.2g/L NaHCO_3._ FBS was reduced to 1.5% in the maintenance medium (MM) which otherwise had similar constituents as CGM.

C6/36 cells were used for propagating dengue serotypes 1-4 in MM at 28°C for seven days. The supernatants were harvested and aliquots were stored at -70°C. Virus titration was done by plaque assay in Vero cells at 37°C with 5% carbon dioxide (CO_2_).

### Cell cytotoxicity assay

Vero 5×10^3^ cells/well in 200 µL of CGM were seeded onto 96-well flat bottomed polystyrene plates (Corning, USA, Cat No. 3598) to obtain 80% confluent monolayers after 24 hours’ incubation at 37°C in 5% CO_2_. Cells were exposed to extract of GG at concentrations ranging from 33.3-1000 µg/mL. Each concentration was tested in triplicate and plates were incubated for seven days at 37°C with 5% CO_2_. Following this the supernatant was removed and 50 µL of 3-(4, 5-dimethyl-2-thiazolyl)-2, 5-diphenyl-2H-tetrazolium bromide (MTT) solution (5 mg/mL) was added to the wells and incubated at 37°C for two hours. The supernatant was then removed and replaced with 100 µL of acidic isopropanol to solubilize the precipitate, and the absorbance was determined at 570 nm and 630 nm (Multiscan microplate spectrophotometer, Model No. 1530, Multisky, Thermo Fisher Scientific, USA). Dose–response curves of cell viability, with reference to the positive and negative controls and log concentrations of the extracts, were plotted using GraphPad Prism software (version 9.0.0.) and the 50% cytotoxic concentration (CC_50_) was calculated. The positive control, chloroquine diphosphate (CQ; Sigma Aldrich, USA), was tested at 0.1-25 µg/mL. Fractions E and F of GG extract were also tested at 2.5-250µg/mL concentrations.

### Virus titration

Virus titration was performed in Vero cells according to the protocol kindly shared by Prof. Damonte, Laboratorio de Virología, Universidad de Buenos Aires, Argentina, with a few modifications. The seeding density of cells and volume of virus inoculum needed to be optimised. Twenty-four well polystyrene plates (UltraCruz, Santa Cruz Biotechnology, USA, Cat No. sc-204445) were seeded with 1.75×10^5^ Vero cells/well in 500 µL of CGM, and incubated at 37°C in 5% CO_2_ atmosphere for 24 hours to obtain 80% confluent monolayers. Tenfold dilutions of virus suspension in MM (250 µL per well) were added, and incubated at 37°C with 5% CO_2_ for one hour with periodic agitation. Following incubation, the virus suspension was replaced with 1mL/well of overlay medium (OM) containing 1% methylcellulose (Sigma, USA) in MM. After seven days at 37°C with 5% CO_2_, cells were fixed with 4% w/v formaldehyde (Sigma, USA). Crystal violet (Himedia, India) with 1% w/v in 10% ethanol (Sigma, USA) was used for staining cells. Each dilution of virus was tested in triplicate wells and two independent titrations were performed. The average plaque count was used to calculate the virus titre and was expressed as plaque forming units per milliliter (PFU/mL).

### Plaque reduction assay to evaluate antiviral activity of GG extract, fractions E and F in Vero cells

Plaque reduction antiviral assay was performed on Vero cell monolayers with all four dengue virus serotypes as described by Talarico et al, except for a few modifications to seeding density of Vero cells and virus inoculum [13]. Vero cells were seeded at 1.75 x10^5^ cells/well in 500 µL of CGM, onto 24-well polystyrene plates (UltraCruz, Santa Cruz Biotechnology, USA, Cat No. sc-204445) and incubated at 37°C in 5% CO_2_ for 24 hours, to obtain 80% confluent cell monolayers. Dengue virus (50 PFU/well) in 250 µL MM was added to each well and incubated for one hour under the same conditions, with periodic agitation. Following virus adsorption, virus inoculum was replaced with one milliliter/ well of OM containing GG extract. Doubling dilutions of GG extract starting with the MNTD (10.41-166.67 µg/mL), were added to each well. Positive control wells with 50 PFU/well of virus were left untreated with extract. Negative control wells were not infected with virus but were treated with GG extract. Plates were incubated for seven days at 37°C with 5% CO_2_. Following which the cells were fixed and stained as described above in *Virus titration* method and the plaques were counted. The assays were performed twice and each concentration of GG extract was tested in duplicate wells. Percentage inhibition of virus was plotted against log of GG concentration using GraphPad Prism (Version 9.0.0.) software for calculation of half maximal inhibitory concentration (IC_50_). Percentage inhibition of virus = (plaque count in positive control - plaque count in test well)/ plaque count in positive control x100. SI was calculated as CC_50_/IC_50_ ratio. Sub-fractions E and F of GG extract (2.5-40 µg/mL) were also tested with all four dengue serotypes.

Interpretation of CC_50_ and IC_50_ values were as follows: CC_50_ <10 µg/mL (very strong cytotoxicity); 10-100 µg/mL (strong cytotoxicity); 100-500 µg/mL (moderate cytotoxicity); and IC_50_ <10 µg/mL (good activity), 10-50 µg/mL (moderate activity), 50-100 µg/mL (low activity) and >100 µg/mL (inactive) [14].

### Plaque reduction assay for fractions 1-7 and sub-fractions A-F

Residues of separated bands from thin layer chromatography were weighed, and used to prepare the overlay medium for the plaque reduction assay. The percentage inhibition per unit weight of fraction was used for comparison of antiviral activity of the seven fractions with dengue virus-1 (DV1). All fractions were tested in duplicate wells. Sub-fractions A-F of fraction-2 were also tested similarly.

### Effect of GG extract, fractions E and F on dengue viral adsorption onto Vero cells

The method described by Talarico et al was followed with a few modifications (seeding cell count and inoculum of virus) to elicit the effect on virus adsorption [13]. Confluent monolayers of Vero cells in 24-well plates were infected with 100 PFU/well of DV1 in the presence of GG extract (166.7 µg/mL), or fraction E (40 µg/mL) or fraction F (40 µg/mL) in separate wells and incubated at 4°C for one hour with periodic agitation. Supernatants with unabsorbed virus were then removed and cells were washed twice with phosphate buffered saline (PB)S followed by addition of OM (1mL) to each well. Virus plaques were counted after seven days of incubation at 37°C with 5% CO_2_ [13].

### Cell cytotoxicity of GG extract on monocytes

Human peripheral blood was provided by healthy donors and buffy coat cells were submitted to density gradient centrifugation (400g for 30 minutes in HiSep™ LSM 1077, HiMedia India) to isolate peripheral blood mononuclear cells (PBMCs). PBMCs were suspended in Roswell Park Memorial Institute (RPMI) 1640 with 10% FBS (RPMI-C) and incubated at 37°C with 5% CO_2_ for 90 min to allow monocyte enrichment [15]. Adherent cells were incubated with different concentrations (60–3000 µg/mL) of GG extract in RPMI with 2% FBS (RPMI-M) in triplicate wells, at 37°C in 5% CO_2_ for 48 hours. After incubation supernatant was replaced with 50 µL of 5 mg/mL MTT and incubated for four hours at 37°C in 5% CO_2_ in the dark. MTT was replaced with 100µl of acidic isopropanol to solubilize the formazan and absorbance was read at 570 nm and 630 nm (Multiscan microplate spectrophotometer, Model No. 1530, Multisky, ThermoFisher Scientific, USA). CC_50_ was calculated as given above for Vero cells.

### Effect of GG extract on NS1 antigen production in monocytes

Following separation of PBMC as described above, cells were suspended in RPMI-C and incubated at 37°C in 5% CO_2_ for 90 min to allow monocyte enrichment. Nonadherent cells were removed by washing with sterile PBS, while adherent cells were detached by mechanical cell harvesting in cold RPMI-C. Recovered adherent cells were suspended in RPMI-C and seeded at 2×10^6^ cells/well in 1 mL of RPMI-C and incubated for 24 hours in 24-well plates [15].

The supernatant was then removed and monocytes were infected with DV1 at multiplicity of infection (MOI) of 4.1 per well. The plate was incubated for two hours at 37°C in 5% CO_2_ [15]. Supernatant of infected wells was replaced with 1 µg/mL or 10 µg/mL of GG extract in RPMI-M, while control wells infected with DV1 were left untreated. Plates were incubated for 48 hours at 37°C in 5% CO_2_. Supernatant was collected from infected GG treated and untreated wells and stored at -20°C. NS1 antigen levels were measured using the Platelia™ Dengue NS1 Antigen kit (Cat No. 72830, Bio-Rad, France) according to the manufacturer’s instructions. Plates were read at 450 nm (Multiscan microplate spectrophotometer, Model No. 1530, Multisky, ThermoFisher Scientific, USA) and NS1 ratios were calculated. Student t-test was used for comparing mean NS1 ratios of treated and untreated cells.

### Gas chromatography of GG

Crude extract of GG was submitted to gas chromatography-mass spectrometry (GCMS) The GG powder was submitted to Agilent Technologies 7890B GC System with a triple-axis detector (Agilant Technologies, USA). Inlet temperature and auxiliary pressure controller temperature were maintained at 250°C and 300°C respectively.

### Bioassay guided fractionation of GG

Preparatory thin layer chromatography (PTLC) technique was carried out using solvent system ethyl acetate (EA: 27227, Sigma, USA): methanol (Me: 32213, Sigma, USA): distilled water (DW: 10:01:01) for the fractionation. Solvent system 1-butanol (24124, Sigma Aldrich): acetic acid (33209, Sigma Aldrich): DW (4:1:5) was used for sub fractionation with PTLC.

Aluminium coated Silica gel 60 F_254_ sheets (1.05554.0007, Supelco, Canada) were used as the stationary phase. PTLC plate was dried well and visualized using anisaldehyde spray reagent and UV light.

### High performance liquid chromatography (HPLC) of GG fractions

The HPLC analysis was carried out on Waters 2535, a quaternary gradient module system with a photodiode array detector and separated by the HPLC column, Rliaant C18 5 μm (4.6 mm x 250 mm column). The mobile phase of 1% acetic acid and 10% methanol in distilled water was supplied in a gradient of 10% to 100% over 30 min at 1 mL/min with an injection volume of 10 µL. Chromatograms of 254 nm were recorded.

### Statistical analysis

Ratio of CC_50_/IC_50_ was used as the Selectivity Index (SI). GraphPad Prism software (Version 9.0.0) was used for plotting dose response curves, nonlinear regression analysis and calculations. Student t-test was used for comparing mean NS1 ratios of GG extract treated and untreated cells.

## Results

GG MNTD and CC_50_ were 166.67 µg/mL and 651.9 µg/mL in Vero cells, whereas CQ MNTD and CC_50_ were 10.0 µg/mL and 17.0 µg/mL respectively (Figure 1).

**Figure 1.**
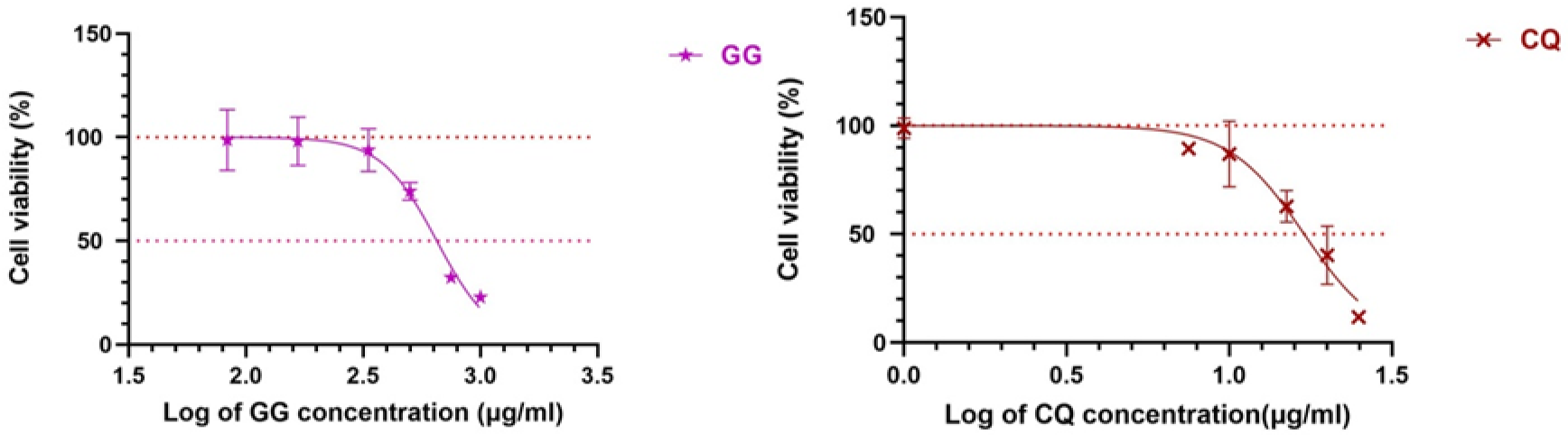
Percentage of Vero cell viability following treatment with different concentrations of (A) aqueous extract of *Glycyrrhiza glabra* (GG) root (33.33-1000 µg/mL) and (B) CQ (0.1-25 µg/mL) incubated with Vero cells for seven days at 37°C in 5% CO_2_. Data points are the mean of triplicate wells ± SEM. Nonlinear regression analysis was performed by GraphPad Prism software to calculate CC_50_.

Table 1 gives the IC_50_, CC_50_ and SI values of GG extract in Vero cells infected with all four dengue serotypes which is graphically represented in Figure 2. GG extract was 98-100% inhibitory to all four serotypes at the MNTD (Figure 2) and had IC_50_ values between 10-50 µg/mL indicating moderate pan-serotype inhibitory activity (Table 1). GG extract had low cytotoxicity for Vero cells as CC_50_ was >500 µg/mL. GG extract SI for all four dengue serotypes were >10, indicating it had good potential as an antiviral agent (Table 1).

**Figure 2.**
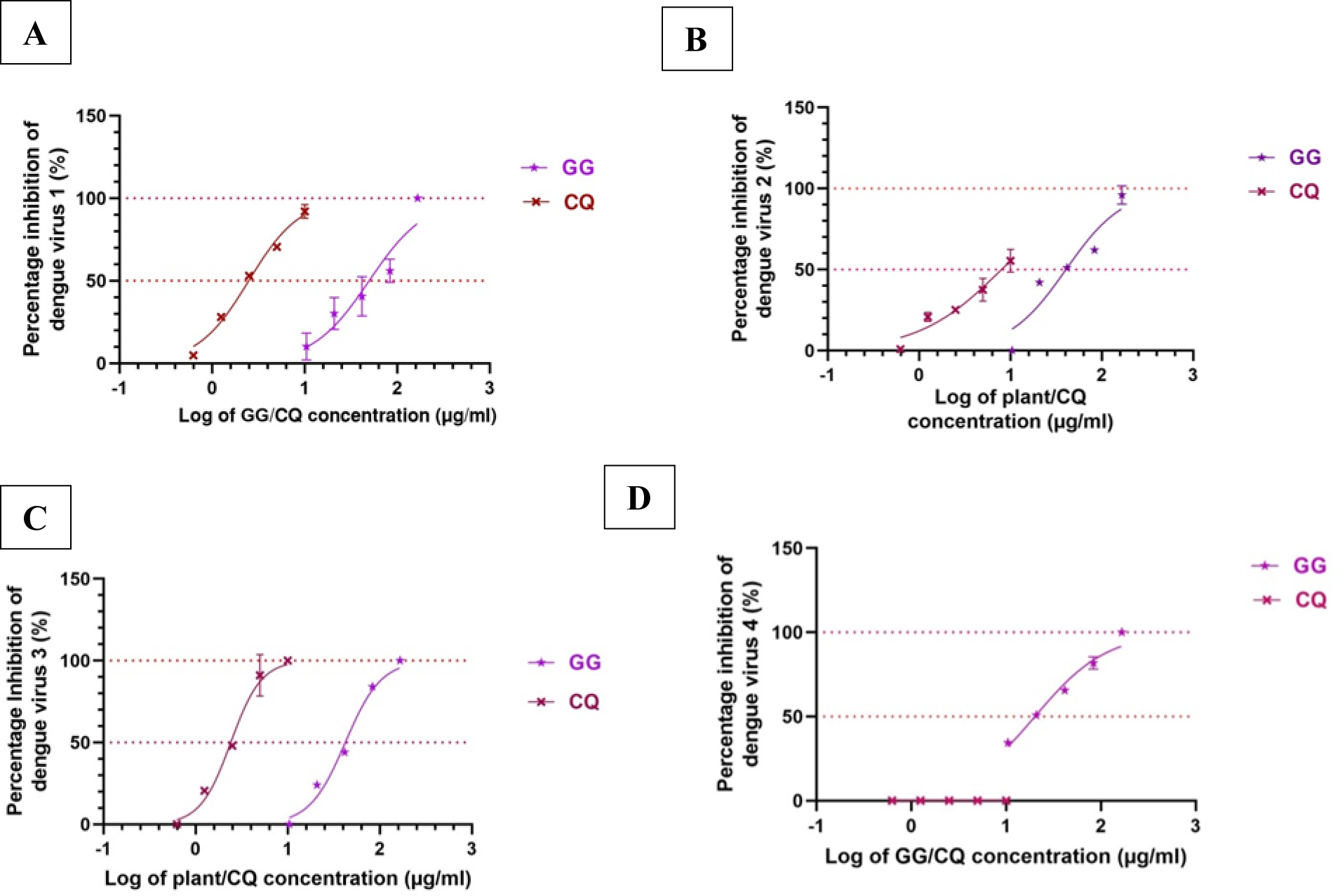
Percentage inhibition of (A) dengue serotype 1 (B) dengue serotype 2 (C) dengue serotype 3 and (D) dengue serotype 4, treated with aqueous extract of *Glycyrrhiza glabra* (GG; 10.4-166.7 µg/mL) and chloroquine diphosphate (CQ; 0.6-10 µg/mL) using plaque reduction assay. Vero cells infected with 50 PFU/mL of virus and incubated for 7 days at 37°C in 5% CO_2_ Each value given is the mean of duplicate assays ± SEM. Nonlinear regression analysis was performed by GraphPad Prism software to calculate IC_50_.

**Table 1.**
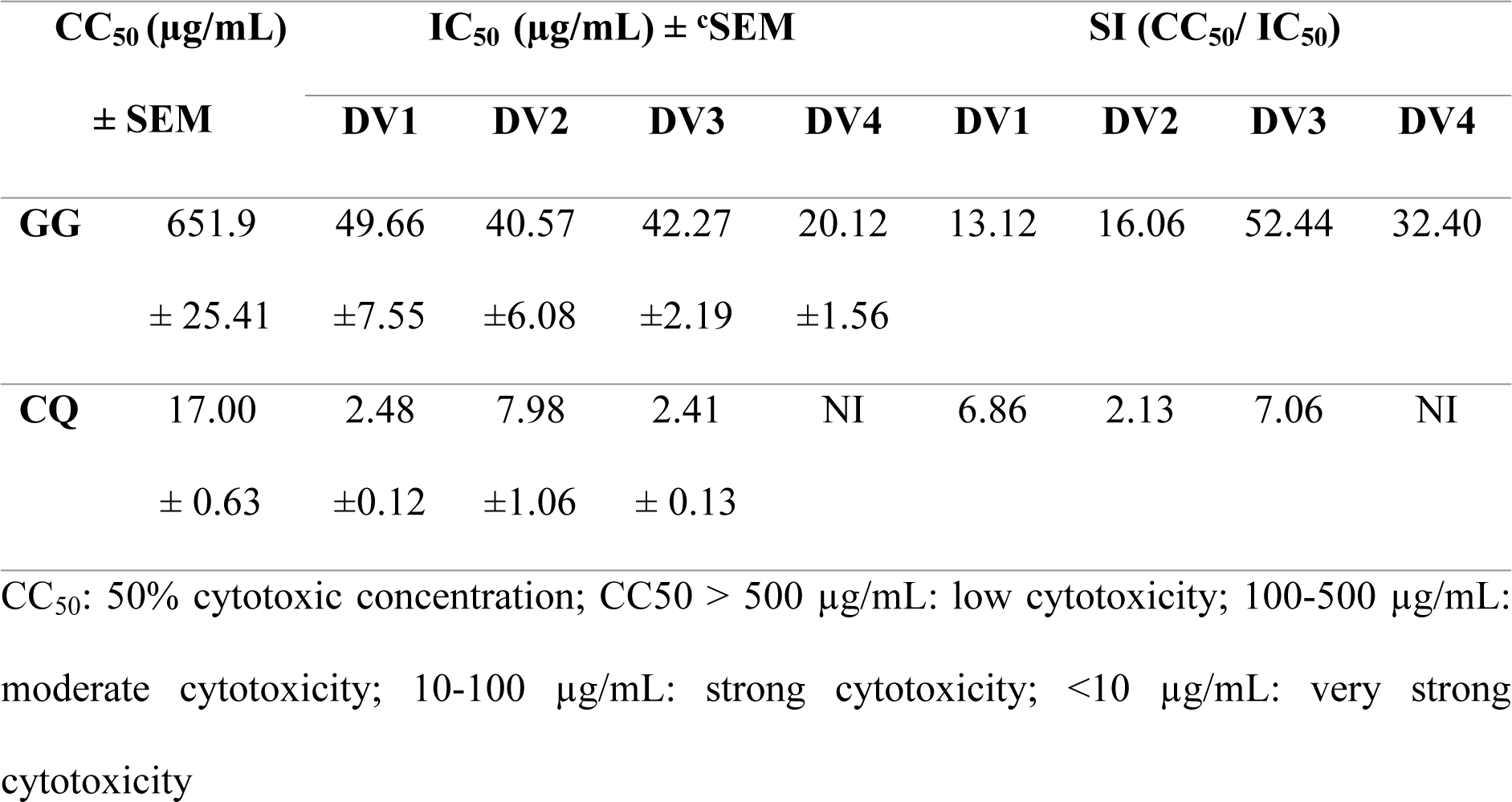
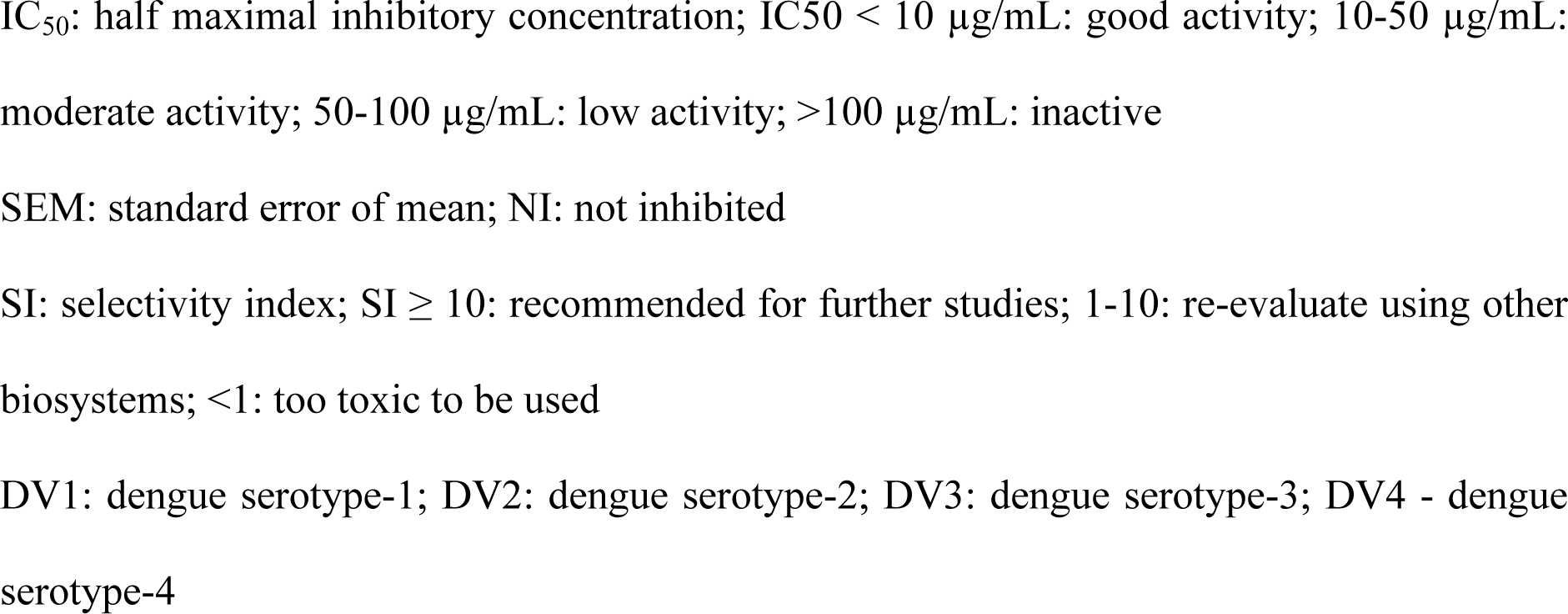
CC_50_, IC_50_ and SI values of *Glycyrrhiza glabra* (GG) and chloroquine (CQ) in Vero cells

| CC<sub>50</sub> (µg/mL) |  | IC<sub>50</sub> (µg/mL) ± SEM |  |  |  | SI (CC<sub>50</sub>/ IC<sub>50</sub>) |  |  |  |
| --- | --- | --- | --- | --- | --- | --- | --- | --- | --- |
|  | ± SEM | DV1 | DV2 | DV3 | DV4 | DV1 | DV2 | DV3 | DV4 |
| **GG** | 651.9 | 49.66 | 40.57 | 42.27 | 20.12 | 13.12 | 16.06 | 52.44 | 32.40 |
|  | ± 25.41 | ±7.55 | ±6.08 | ±2.19 | ±1.56 |  |  |  |  |
| **CQ** | 17.00 | 2.48 | 7.98 | 2.41 | NI | 6.86 | 2.13 | 7.06 | NI |
|  | ± 0.63 | ±0.12 | ±1.06 | ± 0.13 |  |  |  |  |  |

CC<sub>50</sub>: 50% cytotoxic concentration; CC50 > 500 µg/mL: low cytotoxicity; 100-500 µg/mL: moderate cytotoxicity; 10-100 µg/mL: strong cytotoxicity; <10 µg/mL: very strong cytotoxicity
IC<sub>50</sub>: half maximal inhibitory concentration; IC<sub>50</sub> < 10 µg/mL: good activity; 10-50 µg/mL: moderate activity; 50-100 µg/mL: low activity; >100 µg/mL: inactive
SEM: standard error of mean; NI: not inhibited
SI: selectivity index; SI ≥ 10: recommended for further studies; 1-10: re-evaluate using other biosystems; <1: too toxic to be used
DV1: dengue serotype-1; DV2: dengue serotype-2; DV3: dengue serotype-3; DV4 - dengue serotype-4

### Fractionation of GG

Seven fractions (fractions 1-7) were isolated from PTLC. Details of the sub-fractions and percentage inhibition in DV1 infected Vero cells are given in Supplementary file, tables 1 & 2. Fraction 2 had the highest inhibition per unit weight and PTLC yielded seven sub-fractions (A-G; Supplementary file, table 2). Sub-fractions E and F were selected as they had the highest dengue inhibitory activity. These fractions could not be further separated by TLC.

**Table 2.**
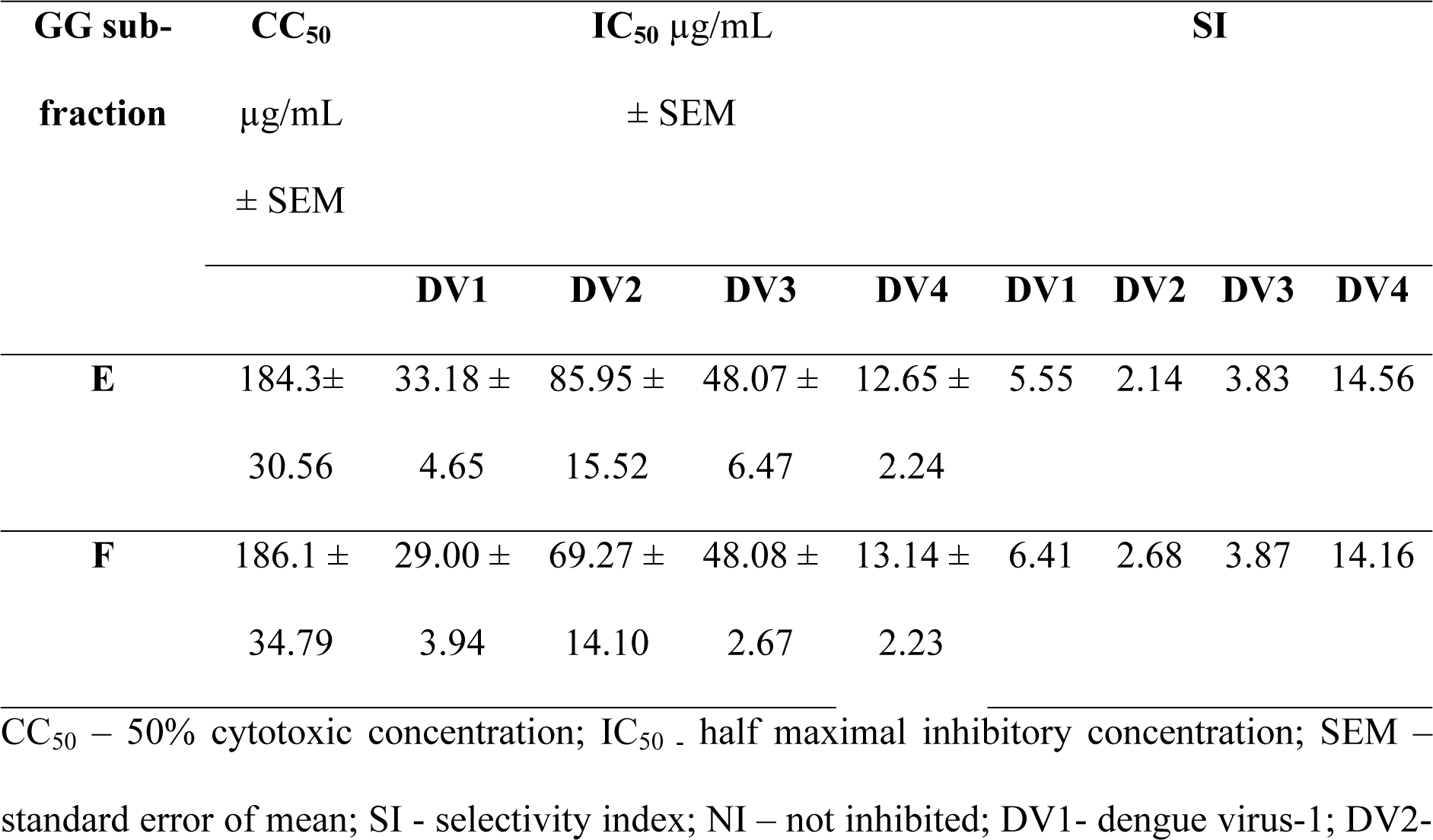
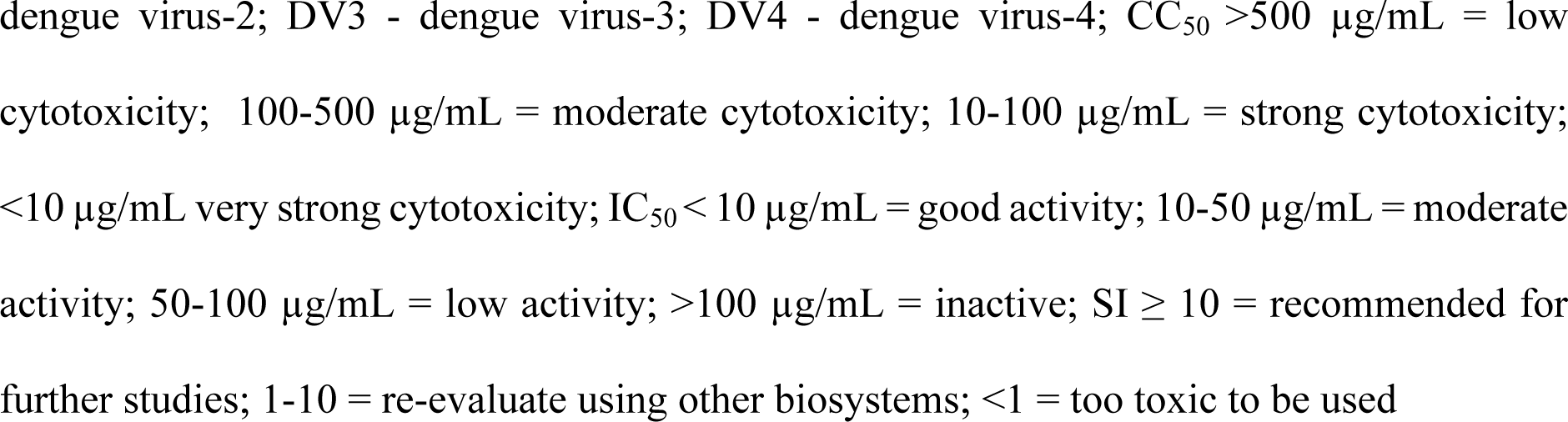
CC_50,_ IC_50_ and SI of GG sub-fractions E and F for dengue serotypes 1-4

### Cytotoxicity of sub-fractions E & F in Vero cells

The MNTD of both sub-fractions were 40 µg/mL and CC_50_ of sub-fractions E and F were 184.3± 30.6 µg/mL and 186.1 ± 34.8 µg/mL respectively (Figure 3, Table 2).

**Figure 3.**
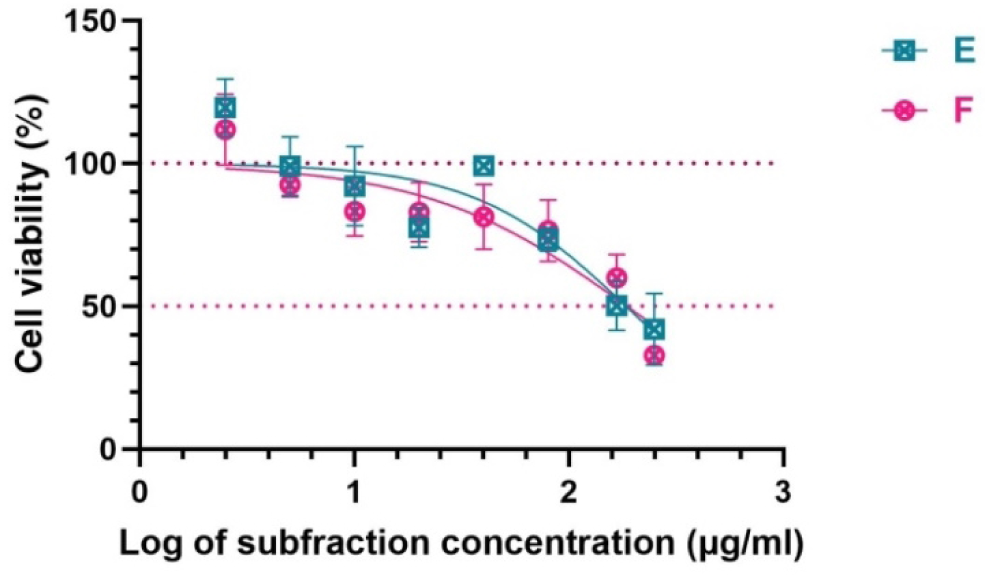
Percentage cell viability following treatment with different concentrations (2.5-250µg/mL) of sub-fractions E and F incubated with Vero cells for seven days at 37°C in 5% CO_2_. Data points are the mean ± SEM of triplicate wells. Nonlinear regression analysis was performed by GraphPad Prism software to calculate CC_50_.

### Inhibition of dengue serotypes 1-4 by sub-fractions E and F

Fractions E and F had low inhibitory activity for DV2 and moderate activity for the other three dengue serotypes; however, only DV4 had a SI > 10 (Table 2, Figure 4).

**Figure 4.**
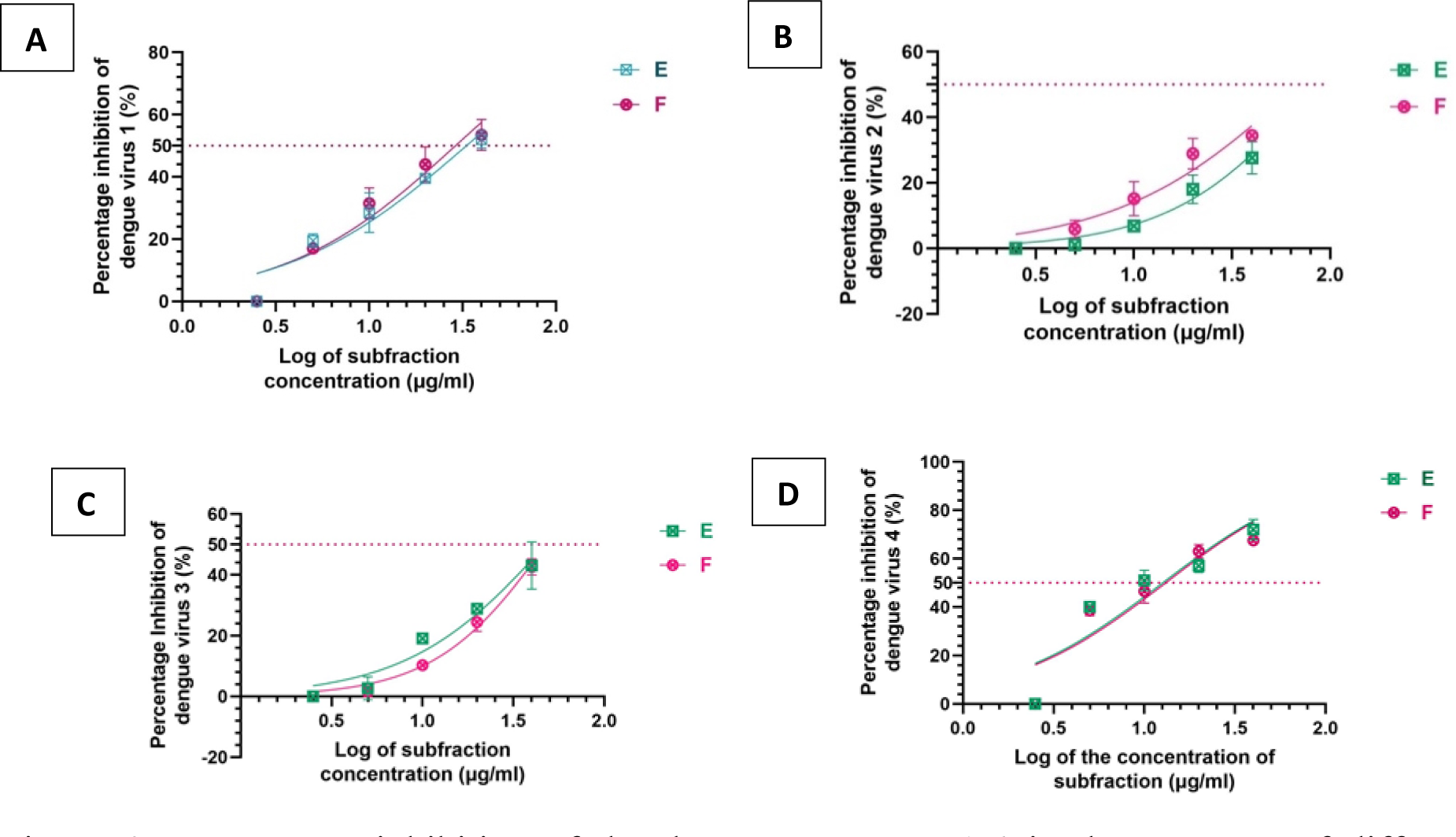
Percentage inhibition of the dengue serotypes 1-4 in the presence of different concentrations (2.5-40 µg/mL) of sub-fractions E and F. (A) dengue virus type-1 (B) dengue virus type-2 (C) dengue virus type-3 (D) dengue virus type-4. Extra-cellular virus yields were determined seven days’ post-infection, by plaque reduction assay in Vero cells infected with 50 PFU/mL of the virus. Each value is given as the mean ± SEM of two independent assays

### Effect of GG extract, fractions E and F on dengue viral adsorption on Vero cells

The effect of GG extract, fractions E and F on adsorption of dengue virus onto Vero cells was ascertained by incubating cells infected with DV1 at 4°C with either GG extract or the fractions. Incubation at 4°C allows adsorption of virus but prevents entry into cells. Crude extract GG (average percentage inhibition 50.4 ± 3.7) and sub-fraction E (average percentage inhibition 24.9 ± 3.2) were capable of reducing virus adsorption of DV1 on Vero cells. Effect of sub-fraction F (mean percentage inhibition 8.3 ± 5.3) was minimal (Figure 5).

**Figure 5.**
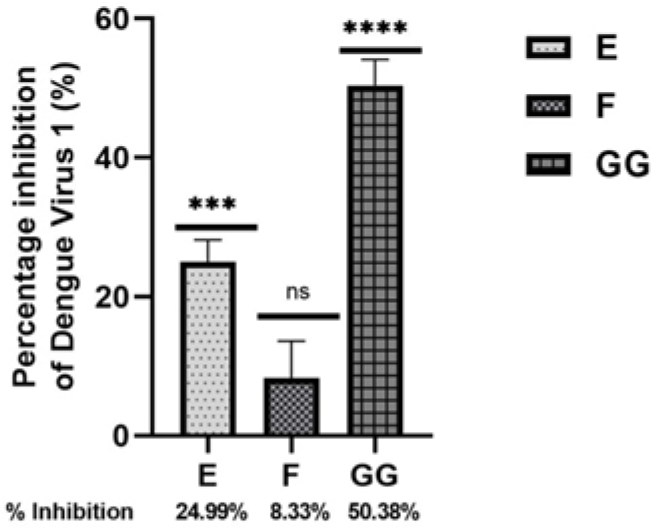
Inhibition of DV1 adsorption by crude extract of *Glycyrrhiza glabra* (GG; 166.7 µg/mL), sub-fractions E and F (40 µg/mL each) as determined by plaque reduction assay with each performed in triplicate and control wells without treatment. Percentage inhibition given is the mean ± SEM of two independent assays, analysed by one-way ANOVA Dunnett’s multiple comparisons test. **** p <0.0001, *** p < 0.001, ns = not significant.

### Effect of GG extract on NS1 antigen production in monocytes

Inhibition of NS1 antigen production by GG extract at 10 µg/ml and 1 µg/ml concentrations was 14.1% and 26.5% respectively compared to that of untreated cells. There was a significant difference (p<0.05) in the reduction of NS1 ratios in the cells treated with 1µg/mL when compared to NS1 ratios of untreated cells.

### Identification of compounds in crude extract of GG

The chromatogram and the list of compounds identified via the equipment library are given in Supplementary file, Table 3, Figure 1 & 2.

## Discussion

Our aim was to discover plant compounds capable of inhibiting dengue viral infection in Vero cells and reducing NS1 antigen production in dengue virus infected monocytes. This is the first report of anti-dengue viral activity of GG aqueous extract on all four dengue serotypes. GG performed better than the positive control CQ which only inhibited DV1-3. Lower viral loads and lower NS1 antigen levels reduce the risk of developing severe dengue [8,12]. MNTD (166.67 µg/mL) of the aqueous extract of GG inhibited all four dengue serotypes in Vero cells at a magnitude of two. It was non-toxic (CC_50_ >500 µg/mL) to Vero cells and had moderate inhibitory activity (IC_50_ 20.12-49.66 µg/mL) with SI >10 for all four dengue serotypes in Vero cells. However, Fraction E and F from GG were moderately toxic (CC_50_ 184.3 and 186.1 µg/mL respectively) and had low inhibitory activity for DV2 with moderate activity for the other three dengue serotypes; good antiviral potential (SI > 10) was found only for DV4.

In the plaque reduction assay, the GG extract and fractions were added following virus adsorption and penetration of Vero cells. In order to ascertain if GG and its fractions specifically inhibit adsorption of the virus to cells, a time of addition assay was conducted. Extract of GG interfered with dengue virus adsorption to Vero cells (50% reduction in plaque formation). Fractions E and F were less efficient at preventing virus adsorption. These findings demonstrate that the aqueous extract of GG contains compounds which are absent in fractions E and F, that make it less toxic as well as more active (acting both before and after penetration of the virus into the cell). The loss of efficacy when crude extracts of plants are fractionated, has been observed in other studies too and is probably due to the loss of synergism among phytochemicals or loss of active compounds or lower MNTD used for assays [16].

A recent study demonstrated that the popular Chinese herbal medicine *Glycyrrhizae Radix* et Rhizome (dry root and rhizome of *Glycyrrhiza uralensis* Fisch., *Glycyrrhiza inflata* Bat., and *Glycyrrhiza glabra* L.) was capable of preventing adsorption of DV2 but was incapable of inhibiting virus cycle after internalization. The same study found that Glycyrrhizic acid, Glycyrrhetnic acid, liquiritigenin, and isoliquiritigenin compounds were individually incapable of significant dengue virus inhibition, which was similar to the reduced inhibition of fractions E and F observed in our study [17]. Glycyrrhizin, a compound found in GG, was studied with DV1, DV2 and DV4 previously and was found to be a potent inhibitor of flaviviruses [18].

Since the crude aqueous extract of GG had better activity than fraction E and F against all four dengue serotypes, further studies on monocytes were carried out with it. Experiments were conducted with DV1 infected monocytes to determine if extract of GG interfered with NS1 antigen production. Following infection of monocytes with DV1, cells were exposed to extract of GG for 48 hours at concentrations of 10 and 1 µg/mL. A significant (p<0.05) reduction in NS1 antigen levels was seen with 1 µg/mL of GG. We chose DV1 for testing NS1 levels as a considerably higher NS1 concentration has been observed with DV1 infection [19]. Mello et al detected the most marked inhibition of NS1 antigen levels at 24 hours with *Uncaria guianensis* leaf extract treated DV2 infected Huh-7 cells [20]. Hence, further testing for effect with all four serotypes, at different incubation intervals, is needed to determine the optimal duration and concentrations for a significant inhibition on NS1 antigen levels with GG.

Water, methanol and ethyl acetate and their mixtures are found to be highly polar solvents which could be used to isolate hydrophilic and polar compounds [21–23]. The n-butanol: acetic acid: distilled water is a polar solvent system which supports the separation of polar compounds [24]. HPLC chromatogram of fraction E showed six identifiable peaks suggesting the presence of six different molecules, while the chromatogram of F demonstrated the presence of one dominant peak suggesting the presence of a single compound with mild impurities. However, both E and F had a common pattern of well-defined peaks at retention times RT;36.673 and RT;36.754 suggesting the presence of closely related compounds in both sub-fractions. However, there were no standards to elucidate the identity of compounds.

The UV spectrum of peaks at 36.678 minutes showed two absorption bands at 228 and 277 nm which are very common in phenolic compounds [25]. However, the absence of UV absorption between 310-350 nm suggested that the above compound is not a flavonoid [26]. Since the preparative TLC fractions, both E and F were soluble in highly polar solvents such as methanol and water, the compounds must contain some polar moieties as well. All this evidence shed some light on the presence of phenolic glycosides in fractions E and F which might contribute to antiviral properties. Since the active fractions are polar, direct injection of sample to GC may not give informative results. Hence pre-column derivation of samples is recommended.

## Conclusion

Aqueous extract of *Glycyrrhiza glabra* roots inhibited all four dengue serotypes significantly and also inhibited NS1 antigen production by dengue virus 1 infected monocytes. *Glycyrrhiza glabra* aqueous extract interfered with adsorption of virus to Vero cells and on virus replication processes after virus internalization. However, fraction E and F were not as successful as the GG aqueous extract in inhibiting all serotypes of dengue. Identification of the compounds that constitute the aqueous extract of *Glycyrrhiza glabra* is needed to produce pharmaceutical agents with antiviral properties.

## Ethical considerations

The Ethics Review Committee of the Faculty of Medicine, Sir John Kotelawala Defence University, approved the application for drawing blood from healthy individuals (RP/2022/04).

## Acknowledgement

Dengue virus serotype 2, Vero and C6/36 cells were provided by Prof. G. N. Malavige, Center for Dengue Research, University of Sri Jayewardenepura, Nugegoda, Sri Lanka. We are grateful to Prof. E.B Damonte, Laboratorio de Virología, Universidad de Buenos Aires, Argentina, for sharing virus titration protocol and Mr. Ranga Tudugala for assistance in statistical analysis.

## Author contributions

KGJ: Investigation, original draft preparation

KG: Funding acquisition, conceptualization, methodology, supervision of cell culture work, project administration, original draft preparation

SS: Supervision of fractionation, original draft preparation

CG: Funding acquisition, supervision of cell cytotoxicity assays, review & editing

PS: supervision of cell cytotoxicity assays, review & editing

NP: performed HPLC

AJ: performed NS1 assays

## Supporting information

S1 Table 1. Retention factor (Rf), yield of the fraction and percentage inhibition of fractions separated from PTLC

S1 Table 2. Retention factor (FR), yield % (w/w), percentage inhibition

S1 Table 3. Compounds identified in the crude extract GG using GCMS

S1 Figure 1. HPLC chromatogram of subfractions E and F

S1 Figure 2. GCMS chromatogram of crude extract GG

